# Towards forecasting antimicrobial resistance evolution: concordance of *in vitro* and *in vivo* studies

**DOI:** 10.64898/2026.10.22.760874

**Authors:** Maren Herzog, Ronan A. Murphy, Jens Rolff

## Abstract

Antimicrobial resistance is a major global health concern, and current measures to address the problem are new drugs, prudent use and rapid diagnostics. Recently, the idea of forecasting resistance evolution experimentally to prevent resistance *a priori* has gained attention. But how well do experimental data on antimicrobial resistance evolution reflect resistance in strains from patients?

We conducted a systematic review and analysis to compare *in vitro* and *in vivo* data. We collected and analysed literature on experimental resistance evolution published between 1977 and 2024 available on PubMed for ESKAPE pathogens plus *E*. *coli* (ESKAPEE). We catalogued the magnitude of resistance evolution via minimum inhibitory concentration (MIC) changes across antimicrobial classes. We then focussed on *P. aeruginosa* and *E. coli* under ciprofloxacin (CIP) selection, as these were most frequent bacterial species used in experimental evolution. CIP-selected gene mutations and amino acid substitutions, as well as the associated MICs in experimentally evolved *P. aeruginosa* and *E. coli* isolates, were compared with corresponding *in vivo* data.

*In vitro* and *in vivo* isolates of *E. coli* showed high concordance in the selected gene mutations and the specific amino-acid changes. Similar results were found for *P. aeruginosa* Evolved MIC values differed significantly depending on the respective gene mutation and the total number of mutations, but not due to the origin of the strains – *in vitro* or *in vivo*. Our findings clearly show that experimental resistance evolution *in vitro* could predict the emergence of antimicrobial resistance evolution *in vivo*.

This study supports the idea that experimental data can be used to forecast the risks of resistance emergence and therefore contribute to antimicrobial stewardship. It will be important to investigate how generalizable these findings are for other bacteria-antimicrobial combinations.

## Introduction

Antimicrobial resistance is a major global health issue, with 5 million deaths associated with AMR in 2019 (Murray, Stanton et al. 2021). Current policies to contain AMR are highly variable across countries (Patel, Harant et al. 2023). In general, prudent use, rapid diagnostics and new drugs are considered to be the main measures to manage and prevent AMR (Zur Wiesch, Kouyos et al. 2011). Over the last few years, the idea of forecasting AMR evolution has been proposed (Martínez, Baquero et al. 2007, Andersson 2015, Rolff, Bonhoeffer et al. 2024). Often forecasting relies on computational methods that use available data for particular drug/disease combinations (Blanquart, Lehtinen et al. 2017, Moradigaravand, Palm et al. 2018, Stracy, Snitser et al. 2022, Aga, Moyo et al. 2026). Such analyses have great potential to manage resistance (sensu Zur Wiesch, Kouyos et al. 2011) that has already emerged, much less so to prevent resistance evolution. An alternative approach is to predict resistance evolution from pharmacodynamics or experimental evolution of resistance (Angst, Tepekule et al. 2021, Rolff, Bonhoeffer et al. 2024). Such predictions would allow management of AMR proactively by prevention rather than managing resistance after it has already emerged (Zur Wiesch, Kouyos et al. 2011, Rolff, Bonhoeffer et al. 2024). This hinges on how closely the results of *in vitro* resistance evolution match the bacterial resistance mechanisms observed in patient or environmental samples: this has not been assessed systematically.

Experimental evolution has been first used in the 19^th^ century (Dallinger 1878) and evolutionary biologists often use this to investigate principles of evolution using antimicrobials as selective agents, a source we can draw on. The ease of controlling for specific levels of stress in microbial cultures is a major strength of *in vitro* experimental evolution, allowing precise links to be drawn between specific culture conditions and changes in populations in response. These observations can be tracked down to the single nucleotide polymorphism (SNP) level in the genome. Yet experimental evolution on AMR is not common practice, for example, during drug development. Experimental evolution has helped illuminate the emergence of resistance to antibiotics; mechanisms of resistance, the conditions under which resistance arises and reservoirs of resistance. Further to this, the linkages drawn between experimental evolution and the molecular mechanisms of resistance using genomics allow for molecular epidemiological tracking of AMR.

Results of experimental evolution experiments may also be vital in predicting and preventing the emergence of resistance in the future. Such experiments provide knowledge of how bacteria react to novel treatments and help to avoid past mistakes of antibiotic use. They can provide opportunities to monitor real-world emergence and spread of resistance and the development of early warning systems about imminent dangers of community resistance. Some studies of particular bacteria compared specific resistance mutations *in vitro* versus *in vivo* and found consistency between experimental and clinical data (for *Mycoplasma hominis*, for *P. aeruginosa* and *Acinetobacter baumannii* see Scribner et al. 2020).

Here we compare the outcome of resistance evolution from experiments with publicly available *in vivo* data and investigate correlations at the amino acid level. We then ask if experimental data can provide useful predictions for the evolution of AMR. We systematically reviewed studies which conducted experimental evolution on bacteria using antimicrobials. From these metadata, we can test for the consistency and robustness of experimental evolution; does a given combination of bacterial species and antibiotic lead to similar outcomes in magnitude of resistance and resistance mutations?

## Methods

### Search strategy and inclusion criteria: in vitro

To compare antimicrobial resistance selected *in vitro* with *in vivo* data, data of each kind were systematically collected. First, papers were collected in which bacteria were experimentally evolved *in vitro* against antimicrobials using PubMed. We focused on medically important pathogens: the ESKAPE group – *Enterococcus faecium, Staphylococcus aureus, Klebsiella pneumoniae, Acinetobacter baumannii, Pseudomonas aeruginosa* and *Enterobacter* species – along with the prominent pathogen *Escherichia coli*. The search was defined using specific search terms and term combinations. The strings were entered in the search window according to the guidelines of the *Advanced Search* in PubMed. To narrow down the literature to *in vitro* data, the search terms “experimental evolution” or “serial passage” were chosen. Publications reporting on the use of antimicrobials were addressed using the search terms “antibiotics”, “antimicrobial peptide” and “host defense peptide”. Finally, the respective bacterial species name was added. The search was conducted on the basis of title and abstract in both British and American spellings.

Studies were only included when they met the following criteria; 1. Data papers published in peer-reviewed journals 2. Written in English 3. Published before January 1, 2025 4. Exposed an ESKAPEE pathogen to an antimicrobial over time 5. The bacterial isolate(s) studied must not have been genetically manipulated in its resistance prior to experimental evolution (i.e. addition or removal of antibiotic resistance mechanisms) 6. Empirically measured the development of resistance to the antimicrobial using minimum inhibitory concentrations (MICs) before and after evolution or fold change in MIC due to *in vitro* exposure.

### Screening: in vitro

The results of each PubMed search were exported as nbib file, and results were imported into EndNote 21 (Clarivate Analytics). Duplicates were removed. The list of unique studies was sorted chronologically, and alternate papers were screened for data extraction (MH and RAM). WebPlotDigitizer 5.2 was used to extract data only available in graphs (Automeris LLC).

In experimental studies involving biological replication, each replicate was recorded as a separate isolate. For each isolate, we recorded the species name, antimicrobial agent, MIC data and, where available, genomic data down to the level of amino acid changes.

### Search strategy and inclusion criteria: in vivo

Most data on experimental resistance evolution were from *P. aeruginosa* and *E. coli* against CIP, these were chosen as our *in vivo* comparators.

Search strings comprised the species name and “ciprofloxacin”, along with resistance genes identified from our *in vitro* data and, where available, amino acid changes in those genes. Inclusion criteria included: 1. Data papers published in peer-reviewed journals 2. Written in English 3. Published before January 1, 2025 4. Studied CIP resistance in *P*. *aeruginosa* or *E*. *coli*5. Studied isolates from an *in vivo* source (e.g. infection, environment, animals) 6. MIC and gene with its amino acid changes distinguishable at the isolate level 7. No *in vitro* manipulation of the isolate prior to MIC determination and sequencing.

### Screening: in vivo

Duplicates were removed for inclusion assessment and data extraction as before (MH and RAM). Along with our inclusion criteria data (species, MIC, amino acid changes), we also recorded categories of the data source (clinical, industrial, environmental, etc.).

### Data analysis

To test whether the origin of the resistant strains – *in vitro* or *in vivo* – had an effect on the logtransformed evolved MIC, we fitted linear mixed effects models for each species. The first model considered the data type and gene mutations as fixed effects; the second model considered the data type and total number of mutations as fixed effects. In both, the reference title was included as a random effect to account for the variability in experimental conditions. Only references were considered in which CIP was used as monotherapy and the MIC values exceeded the EUCAST breakpoint. The first test examined the differences in MIC values between *in vitro* and *in vivo* for all CIP-selected gene mutations shared by both groups. A second test was performed to compare the MIC values between the two groups as a function of the total number of mutations in *P. aeruginosa* and *E. coli* isolates. All data were analysed using RStudio, version 2026.9.0.174 (R Core Team 2025).

## Results

### Composition of data

After screening and data extraction, 62 publications remained (Fig. S1). The resulting publication list was dominated by *P. aeruginosa*, *E. coli* and *S. aureus* (Fig. 1A). The most commonly used antimicrobials in the experimental evolution publications were AMPs, aminoglycosides, fluoroquinolones and beta-lactams (Fig. 1B). Of the individual antimicrobials, CIP was the most common (Fig. 1C). The most commonly tested combinations of bacterial species and antimicrobial were *P. aeruginosa* (8 publications) and *E. coli* (7 publications) evolved with CIP (Fig. 1D).

**Fig. 1.**
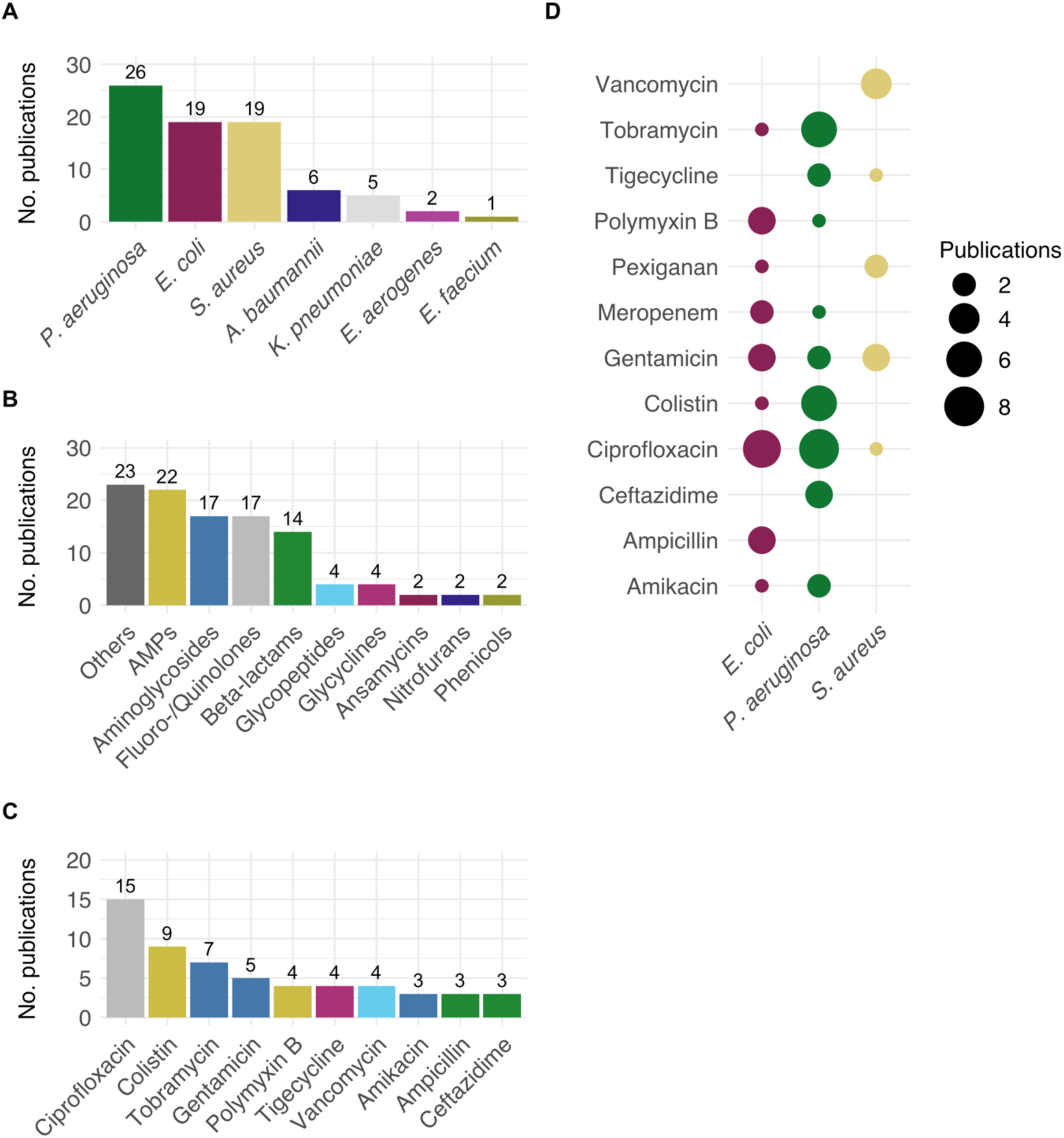
Composition of data resulting from systematic searches and screening for *in vitro* evolution publications. Number of publications featuring each of the ESKAPEE pathogen species (A). Number of publications featuring antimicrobial classes (minimum of 2 occurrences) (B). Number of publications featuring a particular antimicrobial (minimum of 3 occurrences) (C). Number of publications featuring an antimicrobial and its usage with the 3 most common species tested (D).

### Resistance increases due to experimental evolution

We calculated the endpoint fold increase in MIC (evolved MIC/ancestral MIC) for each evolved isolate of each species across our dataset (Fig. 2A, B, C).

**Fig. 2.**
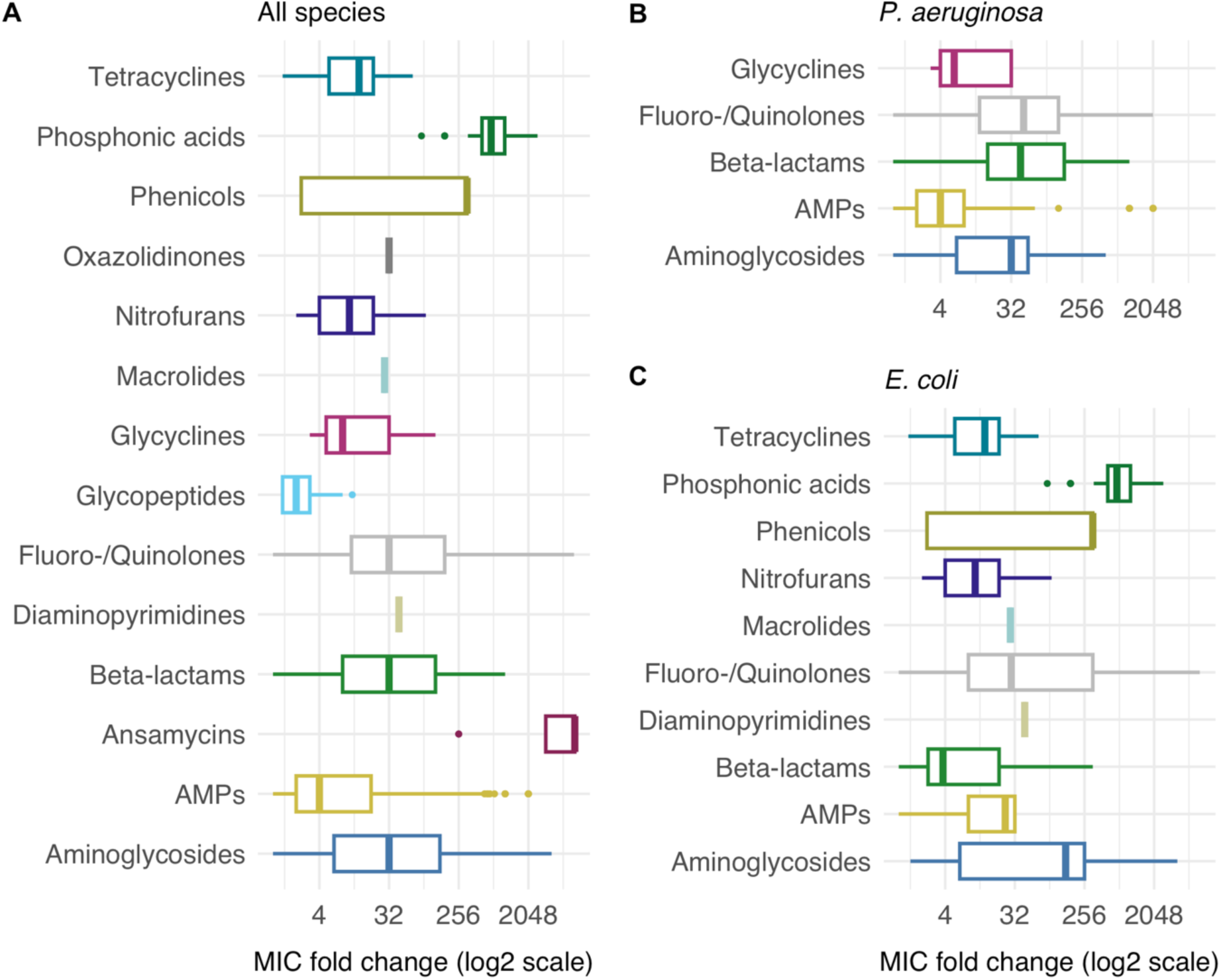
MIC Fold changes within antimicrobial classes across species. Box plots illustrate the extracted fold changes in MIC data resulting from experimental resistance evolution (n = 1,323). A median increase in MIC was observed ranging from 0.25- to 8192-fold across species. This includes findings for all recorded species (A), as well as specific data for *P. aeruginosa* (n = 660) (B) and *E. coli* (n = 423) (C). Antibiotic classes were grouped together into higher-level groups; antibiotic combinations were not considered.

### Mutations Arising During Experimental Evolution

To determine the types and frequencies of mutations resulting from experimental resistance evolution, genomic data from *P. aeruginosa* and *E. coli* were recorded (Fig. 3). We compared resistance mutations against CIP *in vitro* from those found in our *in vivo* searches (Fig 4). Illustrated are the proportions of the five most frequently identified gene mutations in both *in vitro* and *in vivo* isolates of *P. aeruginosa* and *E. coli* (Fig. 5A-D). In *P. aeruginosa* evolved *in vitro* (n = 82), the highest proportion of isolates showed mutations in *gyrA*, followed by *nfxB*, *parC*, *gyrB*, and *rnfC* (Fig. 4A). Analysis of *in vivo*-evolved *P. aeruginosa* isolates (n = 478) revealed an overlap in four out of the five most abundant gene mutations (Fig. 5B).

**Fig. 3.**
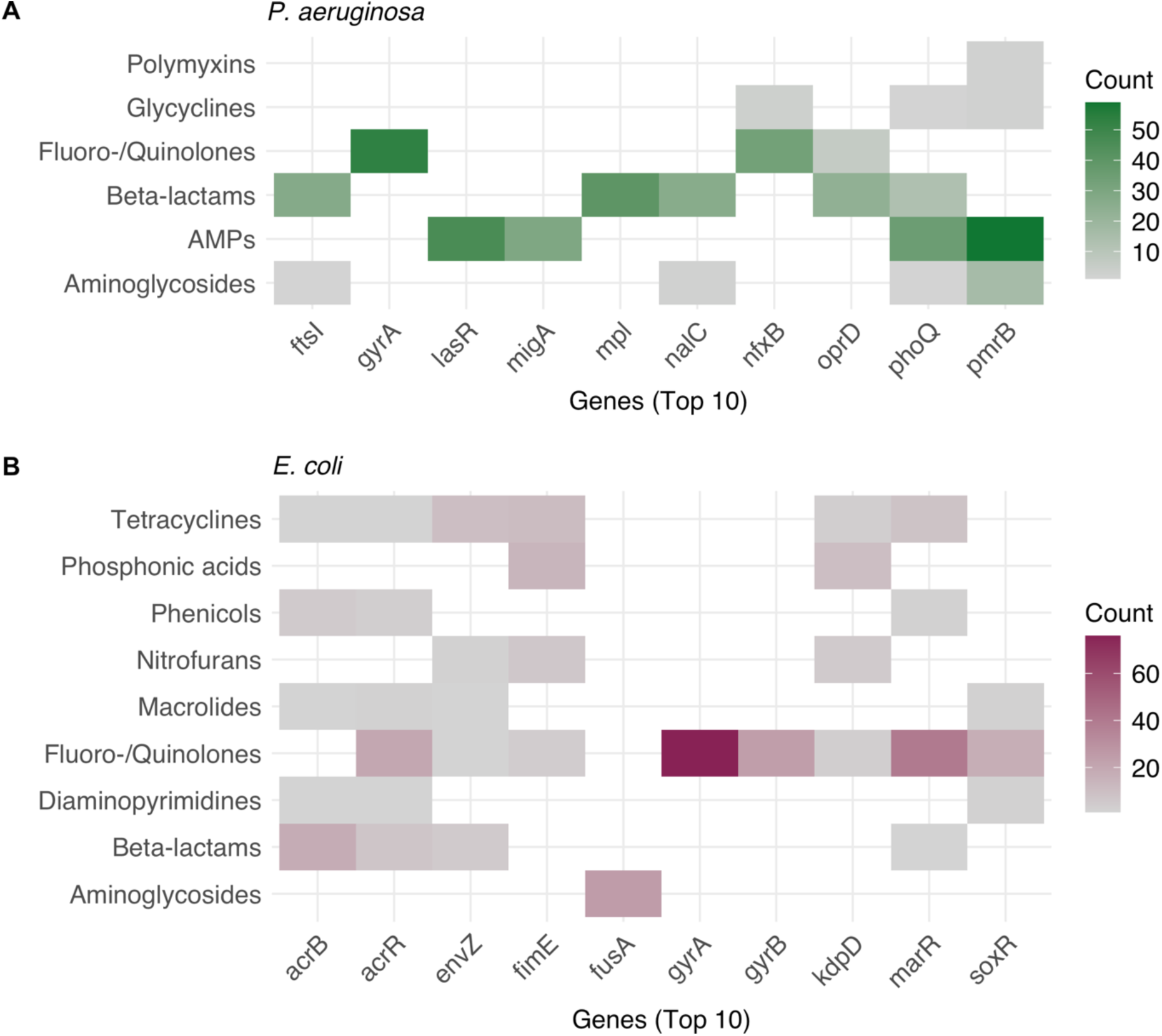
Most frequent gene mutations in *in vitro* isolates across antimicrobial classes. Heat maps show the counts of the ten most frequently reported gene mutations within antimicrobial classes for *in vitro* isolates of *P. aeruginosa* (n = 282) (A) and *E. coli* (n = 247) (B). The highest counts are shown in green (*P. aeruginosa*) and burgundy red (*E. coli*), the lowest are shown in grey. Antibiotic classes were grouped together into higher-level groups; antibiotic combinations were not considered.

**Fig. 4.**
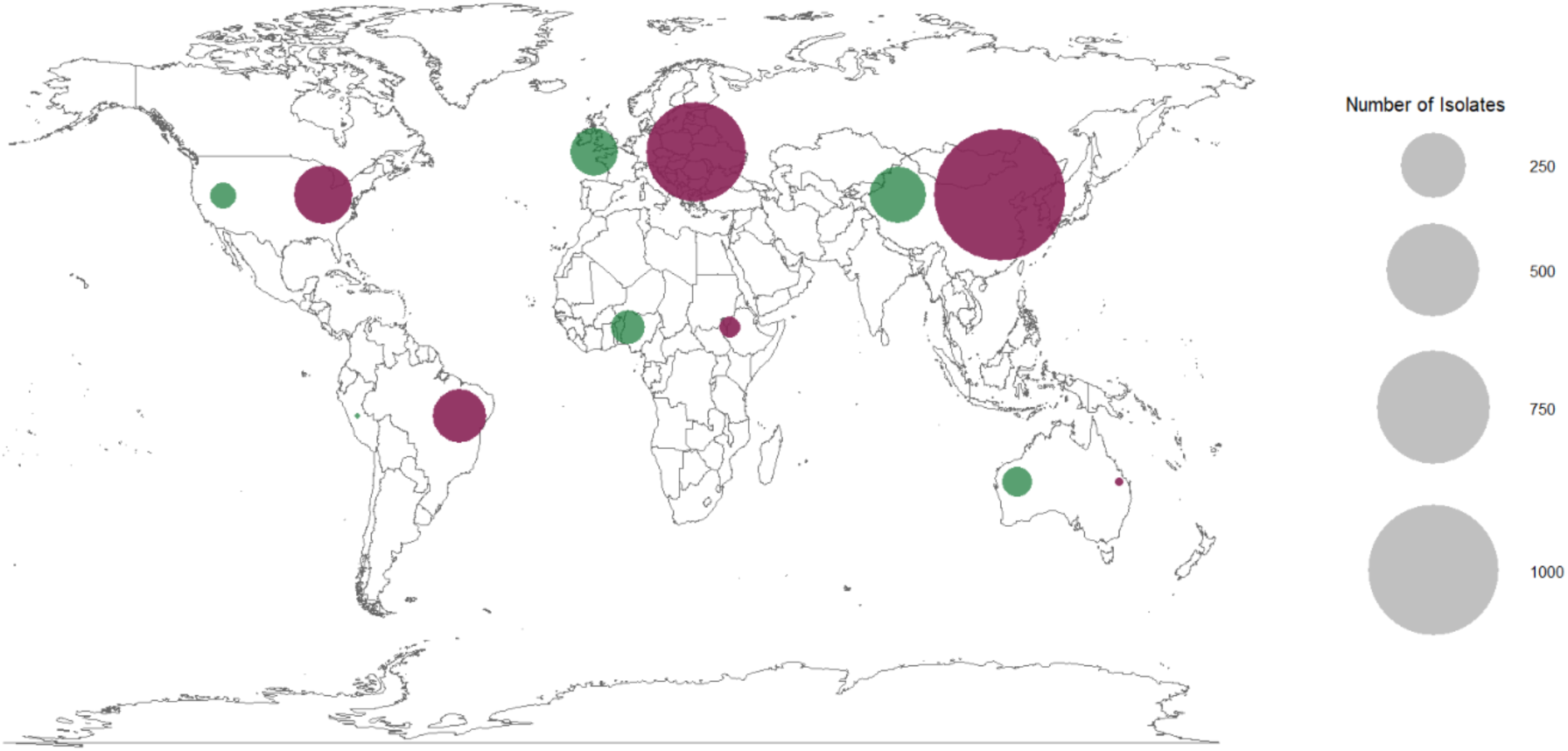
Global sources of *in vivo* isolates of *P. aeruginosa* and *E. coli* included in this study. Circles represent the number of *in vivo* isolates for a given continent for each species. Green circles show *P. aeruginosa,* while burgundy red circles show *E. coli*.

**Fig. 5.**
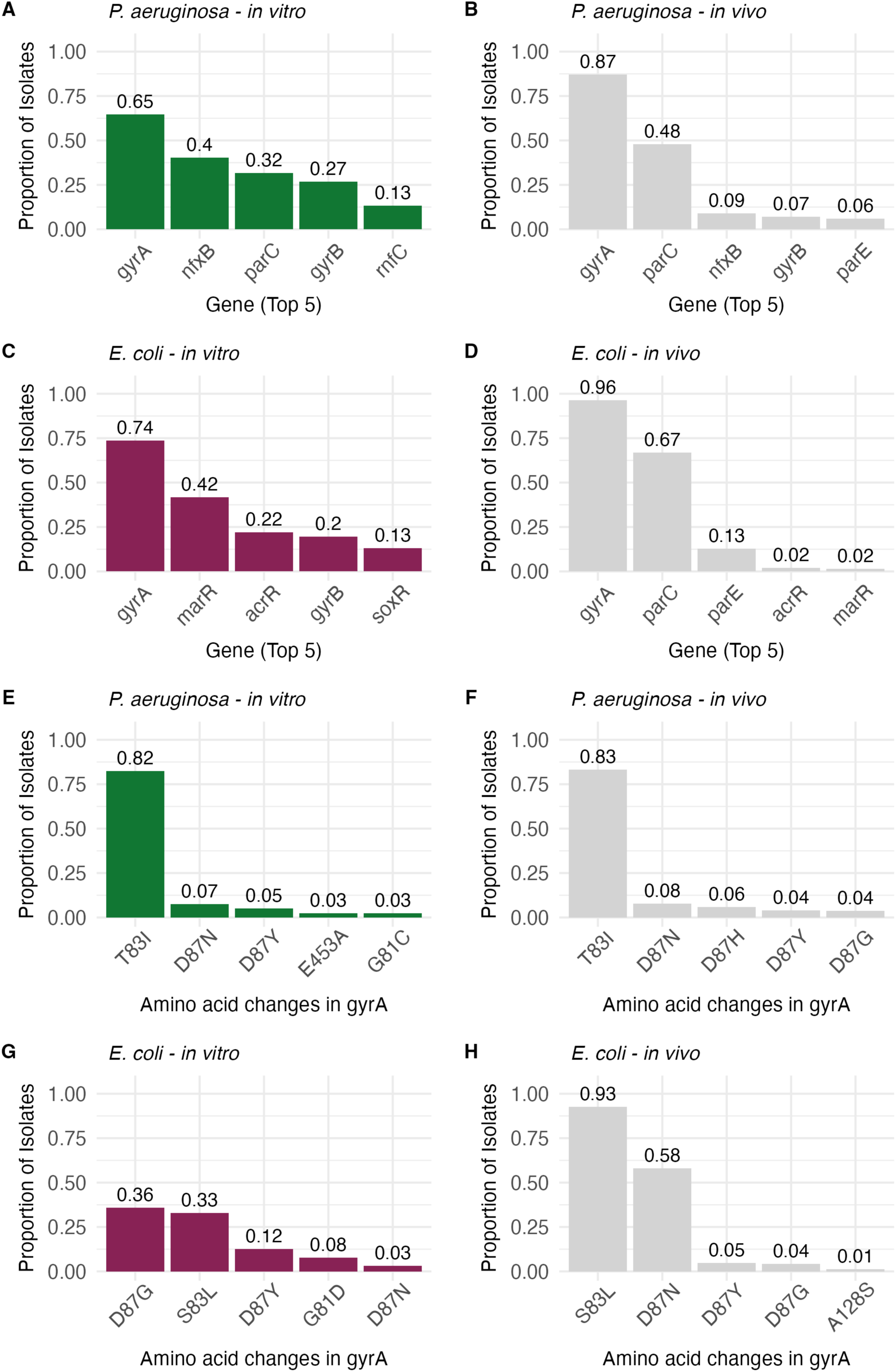
Proportions of gene mutations and amino acid changes underlying CIP resistance. Bars show the proportions of the five most frequently reported gene mutations in the context of CIP resistance in *P. aeruginosa in vitro* (n = 82) (A) and *in vivo* (n = 478) (B), as well as in *E. coli in vitro* (n = 122) (C) and *in vivo* (n = 2,259) (D). The lower part of the panel shows proportions of the corresponding top 5 amino acid changes that cause the mutation in *gyrA* in *P. aeruginosa* evolved in *vitro* (n = 40) (E) and *in vivo* (n = 416) (F) as well as *E. coli* evolved *in vitro* (n = 64) (G) and *in vivo* (n = 2,179) (H). The colour scheme for the *in vitro* data of *P. aeruginosa* (green) and *E. coli* (burgundy red) was retained. *In vivo* data are shown in grey.

A comparison of the five most prevalent gene mutations in *E. coli* isolates from *in vitro* and *in vivo* studies showed concordance for three of the top five gene mutations. The most frequently observed mutations among *in vitro* isolates (n = 122) were identified in *gyrA*, followed by *marR*, *acrR*, *gyrB*, and *soxR* (Fig. 5C). In *E. coli* isolates evolved *in vivo* (n = 2,259), mutations also occurred most frequently in *gyrA* (Fig. 5D).

For a more detailed comparison of *in vitro* and *in vivo* resistance, we looked at the amino acid alterations associated with CIP-selected mutations in *gyrA*. The proportions of the five most prevalent amino acid changes in CIP-resistant *P. aeruginosa* and *E. coli* isolates that harbour mutations in *gyrA* are shown in Fig 5E-H.

To test whether MICs differed between *in vitro* and *in vivo* isolates, we compared MICs across shared resistance-conferring mutations and examined the relationship between MIC and the total number of mutations associated with CIP resistance in *P. aeruginosa* and *E. coli* isolates (Fig. 6).

**Fig. 6.**
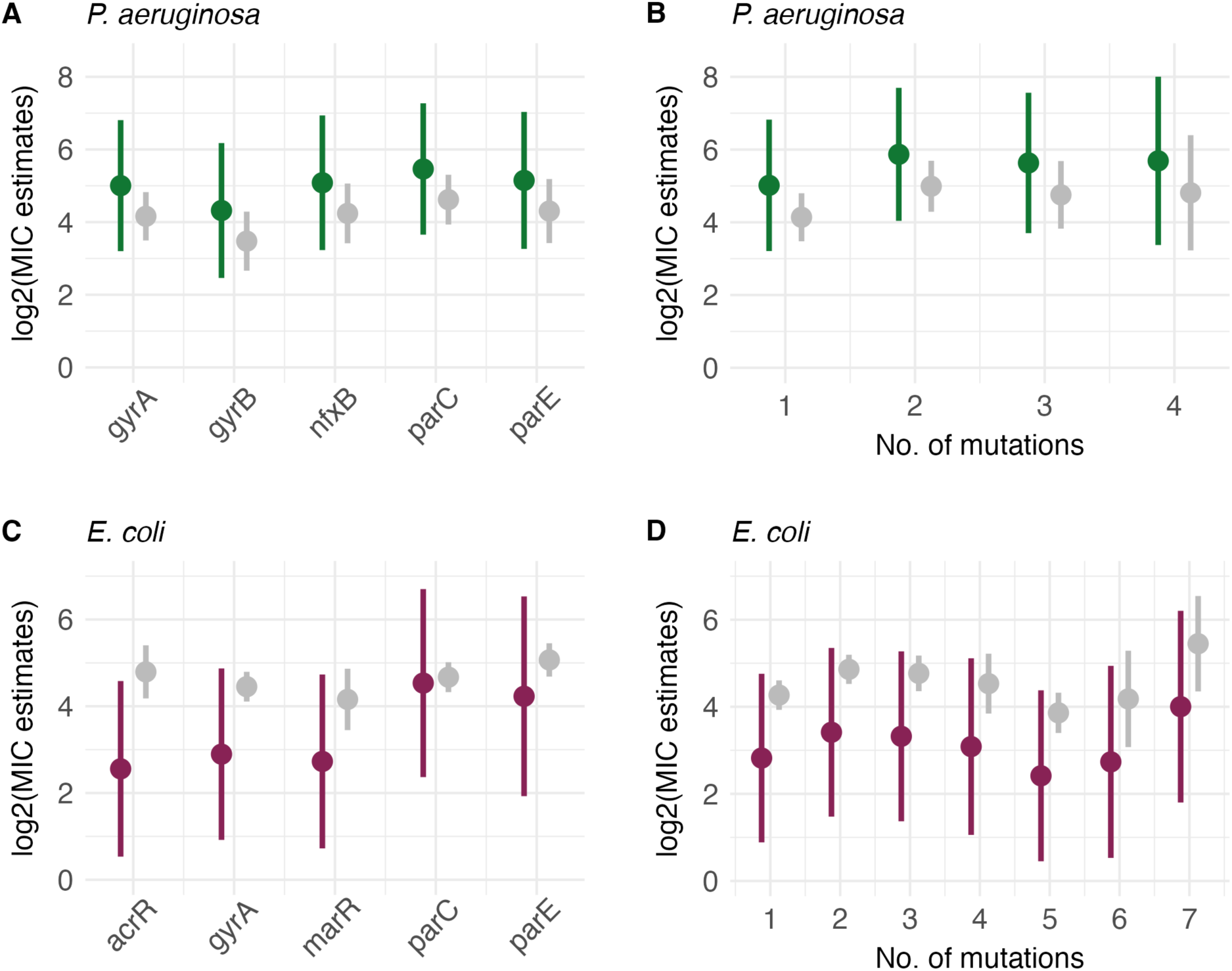
MIC estimates by gene mutation and number of mutations in response to CIP exposure. The box plots show evolved MIC estimates after CIP exposure for the five most frequently selected gene mutations observed in resistant isolates of *P. aeruginosa* (A) and *E. coli* (C) across *in vitro* and *in vivo* studies. Furthermore, the evolved MIC estimates are presented alongside the corresponding number of mutations that emerged from CIP exposure in *P. aeruginosa* (B) and *E. coli* (D) isolates, both *in vitro* and *in vivo*. MIC estimates are log2-transformed. The colour scheme for the *in vitro* data of *P. aeruginosa* (green) and *E. coli* (burgundy red) has been retained. *In vivo* data are shown in grey.

In *P. aeruginosa*, including data type did not significantly improve model fit, either after accounting for the identity of the resistance-conferring mutation (*χ²*_1_ = 0.78, *P* = 0.378) (Fig. 6A) or after accounting for the total number of mutations per isolate (*χ²*_1_ = 0.79, *P* = 0.373) (Fig. 5B). However, MICs varied significantly among specific resistance-conferring mutations (*F*_5/815.78_ = 5.07, *P* < 0.001) and according to the total number of mutations per isolate (*F*_1/935.22_ = 26.85, *P* < 0.001).

In *E. coli*, no significant overall difference in the evolved MIC was observed between *in vitro* and *in vivo* findings across gene mutations (*F*_1/110.0_ = 1.13, *P* = 0.29) (Fig. 6C). However, a significant interaction indicated that the relationship between gene and evolved MIC is not consistent across *in vitro* and *in vivo* findings (*χ²*_8_ = 15.74, *P* = 0.046). A detailed post-hoc analysis indicated that mutations in *acrR* (*P* = 0.04) were primarily responsible for driving this interaction (Table S1). The comparison between *in vitro* and *in vivo* MICs as a function of total mutation count provided no evidence that including data type in the model improved model fit (*χ²*_1_ = 2.6, *P* = 0.107) (Fig. 6D; Fig. 6H). Consistent with prior observations in *P. aeruginosa*, MIC values in *E. coli* varied significantly by gene mutation (*F*_8/3621.8_ = 3.95, *P* < 0.001) and total mutation count (*F*_1/5737.6_ = 79.7, *P* < 0.001).

## Discussion

We have performed the first systematic investigation comparing *in vitro* experimental evolution studies on antimicrobial resistance with real-world resistance. We find that experimentally evolved *in vitro* resistance closely mirrors resistance patterns seen *in vivo*. Focusing our analyses on CIP resistance in *P. aeruginosa* and *E. coli,* we found hardly any differences in CIP MICs with respect to resistance genotype or resistance mutation accumulation between *in vitro* and *in vivo* isolates. These results support the idea that experimentally obtained data are concordant with real-world resistance and have great potential for predicting *in vivo* resistance. The highest proportions of CIP-resistant *P. aeruginosa* and *E. coli* isolates carried mutations in *gyrA* both *in vitro* and *in vivo*. Mutations in *gyrA* and *parC* have previously been documented as the most common CIP target mutations in both *P. aeruginosa* and *E. coli* and they have minimal fitness costs, helping to explain why the same genetic variants are repeatedly selected across *in vitro* and *in vivo* datasets (Marcusson, Frimodt-Møller et al. 2009, Bruchmann, Dötsch et al. 2013, Van Der Putten, Remondini et al. 2019). Within the most frequently mutated gene, *gyrA,* we saw high concordance between *in vitro* and *in vivo* isolates across both species. In both *P. aeruginosa* and *E. coli*, *gyrA* amino acid changes were most frequently observed at positions 83 and 87, as in previous reports (Bruchmann, Dötsch et al. 2013, Van Der Putten, Remondini et al. 2019).

In contrast, mutations affecting drug efflux are closely linked to reduced fitness (Marcusson, Frimodt-Møller et al. 2009). Such constraints can help explain why the same genetic variants are repeatedly selected across *in vitro* and *in vivo* datasets.

While we have developed a unique dataset, it is not without limitations. We saw that genotyping methodologies differed across papers, with whole-genome sequencing common *in vitro* but direct gene amplification predominant *in vivo*. This may bias the apparent frequency of resistance-associated gene mutations, as targeted sequencing in *in vivo* studies may underrepresent mutations in genes not selected *a priori*, unlike whole-genome sequencing. Due to its very limited presence in *in vitro* data, our analysis also does not account for resistance genes acquired via horizontal gene transfer (but see Muetter et al. 2024), which can significantly contribute to the evolution of antimicrobial resistance (Martinez and Baquero 2000). Consequently, the contributions of mobile genetic elements, plasmid-mediated resistance, and gene acquisition could not be systematically compared between *in vitro* and *in vivo*. In addition, while we analysed the relationship between mutation accumulation and MIC, we did not account for the specific combination effects of mutations. The relationship between resistance-conferring mutations and MIC is therefore likely to be simplified, because epistatic interactions among mutations can alter both the magnitude and trajectory of resistance evolution (Wong 2017).

For *in vitro* data, a much shallower pool of genetic backgrounds was present, dominated by a handful of strains. The good match of resistance phenotypes and genotypes across *in vitro* and *in vivo* studies for both *E. coli* and *P. aeruginosa* suggests that the effect of resistance mutations is not strongly dependent on genetic background.

There is likely to be large heterogeneity in the bacterial environments across our dataset. In laboratory settings, sampling intervals, antimicrobial concentrations, temperature, and population size are typically standardised or held constant, whereas these parameters fluctuate substantially within host environments. Bacterial population density is known to affect not only antimicrobial susceptibility (Loffredo, Savini et al. 2021) but also the emergence rates and mechanisms of resistance (Vega and Gore 2014). In experimental evolution, monocultures are used, whereas bacterial populations within the host exist in heterogeneous multi-species communities. Treatment duration impacts the evolution of resistance, with short-term antibiotic exposure favouring low-cost resistance (Yaffe, Dethlefsen et al. 2025). Resistance can evolve from very low concentrations commonly encountered in natural environments (Gullberg, Cao et al. 2011) to high concentrations used therapeutically (Kümmerer 2009). Despite all these sources of variation, our data show that *in vitro* evolution closely recapitulates *in vivo* resistance evolution.

Specific studies have already been performed to test relationships between *in vitro* and *in vivo* antimicrobial resistance. An artificial sputum model designed to mimic lung infection captured the evolutionary diversification of *P. aeruginosa* and produced phenotypes that closely resembled those observed in clinical isolates in cystic fibrosis (Davies, James et al. 2017). In a study by (Jahn, Munck et al. 2017), *E. coli* populations were evolved under four antibiotic selection regimes: antibiotic-resistance-associated mutations and phenotypic shifts emerged consistently across regimes. Our results add a broad evidentiary base in support of these context-specific studies that experimental *in vitro* evolution can reproduce clinically relevant resistance trajectories.

A recent large-scale study of global *Klebsiella* pneumoniae genomes shows that resistance evolution in that species is both consistent and predictable (Aga, Moyo et al. 2026). MICs have already been used in mathematical models that predict the evolution of resistance (Yasir, Karim et al. 2022) and in time-series modelling to predict the future burden of antimicrobial resistance (Kim, Rupasinghe et al. 2023). The data presented here support the idea that resistance evolution can be predicted, as in lab strains under laboratory conditions in our study we found the same resistance outcomes as in a global set of isolates from a variety of sources.

We observed a species-dependent trend in MICs, whereby *P. aeruginosa* isolates obtained from *in vitro* selection showed higher MICs than isolates derived from patients (Fig. 5A), while the reverse was true for *E. coli* (Fig. 5C). This species-specific pattern held true for all gene mutations that occurred in both groups, regardless of the number of mutations per isolate. *P. aeruginosa* is intrinsically less susceptible to many antimicrobial agents than *E. coli*. Its larger genome size, approximately 5.5–7 Mbp compared with 4.6 Mbp in *E. coli* (Pang, Raudonis et al. 2019), may provide broader intrinsic resistance and stress-response capacity (Langendonk, Neill et al. 2021). Consequently, *E. coli* may rely more on accumulating resistance-conferring mutations during antimicrobial exposure (Balachandran, Chatterjee et al. 2025). Ecological differences may further contribute to this pattern. *E. coli* is a commensal in the human gut, whereas *P. aeruginosa* is not among the dominant core residents in healthy individuals. Because antimicrobials perturb the gut microbiota, *E. coli* can experience more frequent collateral selection during therapy. Antibiotic resistance genes in the adult human gut microbiota have been reported to be predominantly associated with tetracycline and fluoroquinolone resistance and to originate largely from *E. coli* (Li, Brejnrod et al. 2023). These species-specific differences are important for resistance forecasting, and predictive frameworks should incorporate species identity, ecological niche, and intrinsic resistance capacity.

One of the few studies directly comparing experimentally selected and *in vivo* resistance tested the mutant selection window for fosfomycin in *P. aeruginosa* and *E. coli* (Pan, Mei et al. 2017). *In vitro*, fosfomycin exposure selected resistant mutants with increased MICs, whereas *in vivo* resistance was species- and regimen-dependent: *P. aeruginosa* evolved resistance under certain dosing conditions, whereas no resistance emerged in *E. coli*. This illustrates an important limitation of experimental evolution studies: simple in *vitro* systems may capture the mutational routes and phenotypic consequences of resistance once it evolves, but they may be less informative about the probability and timing of resistance emergence in complex host environments.

In experimental lab work, replicates of *in vitro* isolates are randomly selected, whereas *in vivo* datasets often rely on single isolates obtained from already-infected patients. Main drivers of AMR are antibiotic use, hospitalisation, surgery, and patients’ clinical histories (Chatterjee, Modarai et al. 2018). These risk factors strongly shape the evolution of antimicrobial resistance in clinical settings, resulting in unequal selection pressures between *in vivo* and *in vitro* settings. Nevertheless, our results show strong concordance between experimentally selected and clinically observed resistant genotypes and phenotypes. This suggests that such clinical risk factors may primarily influence the probability or rate at which resistance emerges, rather than the mutational routes or resistance phenotypes once resistance has evolved (Witzany, Bonhoeffer et al. 2020, Stennett, Back et al. 2022, Garlasco, Arieti et al. 2026). However, because our dataset contains substantially more in vivo than *in vitro* isolates, we cannot resolve this quantitative effect.

### Predict resistance evolution in the future

Our findings show that *in vitro* resistance evolution can closely mirror *in vivo* resistance patterns, supporting its value for forecasting resistance trajectories before they become clinically established. However, translating this potential into robust forecasting will require standardised approaches. For experimental data to support reliable predictions of antimicrobial resistance in the future, experimental conditions should be aligned as closely as possible with clinically relevant settings. Time to resistance and its prevalence may not be accurately captured in simple *in vitro* experiments, as they depend heavily on factors such as the host’s fitness costs and its immune response (Pan, Mei et al. 2017, Pan, Dong et al. 2017). Microfluidic devices designed to mimic natural bacterial niches provide one promising route towards more realistic experimental environments (Zhang, Lambert et al. 2011). Drug-pathogen combinations should be prioritised according to their clinical relevance, and forecasting models should explicitly incorporate species-specific biology and ecological context. Emerging machine-learning approaches that integrate phenotypic and genomic data to predict antimicrobial resistance show promising performance (Adeyemi and Paudel 2026). Linking experimental evolution with clinical resistance dynamics will also require platforms that capture epidemiological processes. Angst et al. used a robotic liquid-handling system that incorporates treatment, transmission, and ‘patient’ admission and discharge, thereby more closely approximating the evolution of resistance in hospital settings (Angst, Tepekule et al. 2021). In a follow-up paper, Muetter et al. 2024 also demonstrated that this approach can be used for resistance acquired by horizontal gene transfer (Muetter, Angst et al. 2024).

Together, these findings support the notion that experimental data can improve the management of antimicrobial resistance by enabling proactive prevention. Taken together with our observation that experimental *in vitro* evolution closely mirrors *in vivo* resistance patterns, these findings suggest that resistance forecasting is feasible using experimental data and could help prevent *de novo* resistance evolution.

## Contributors

Conceptualization: MH, RAM, JR

Data curation: MH, RAM

Formal analysis: MH

Funding acquisition: JR

Investigation: MH, RAM, JR

Methodology: MH, RAM, JR

Project administration: JR

Resources: JR

Supervision: JR

Visualization: MH, RAM

Writing – original draft: MH, RAM

Writing – review & editing: MH, RAM, JR

## Acknowledgements

We thank Mohammed Elbediwi for helping us get off to a good start. We also thank Ofer Zelichover for his invaluable assistance in preparing the data for analysis. We thank Caroline Zanchi for her bioinformatics expertise. We also thank Dino McMahon for comments on the manuscript.

## Supplement

**Fig. S1.**
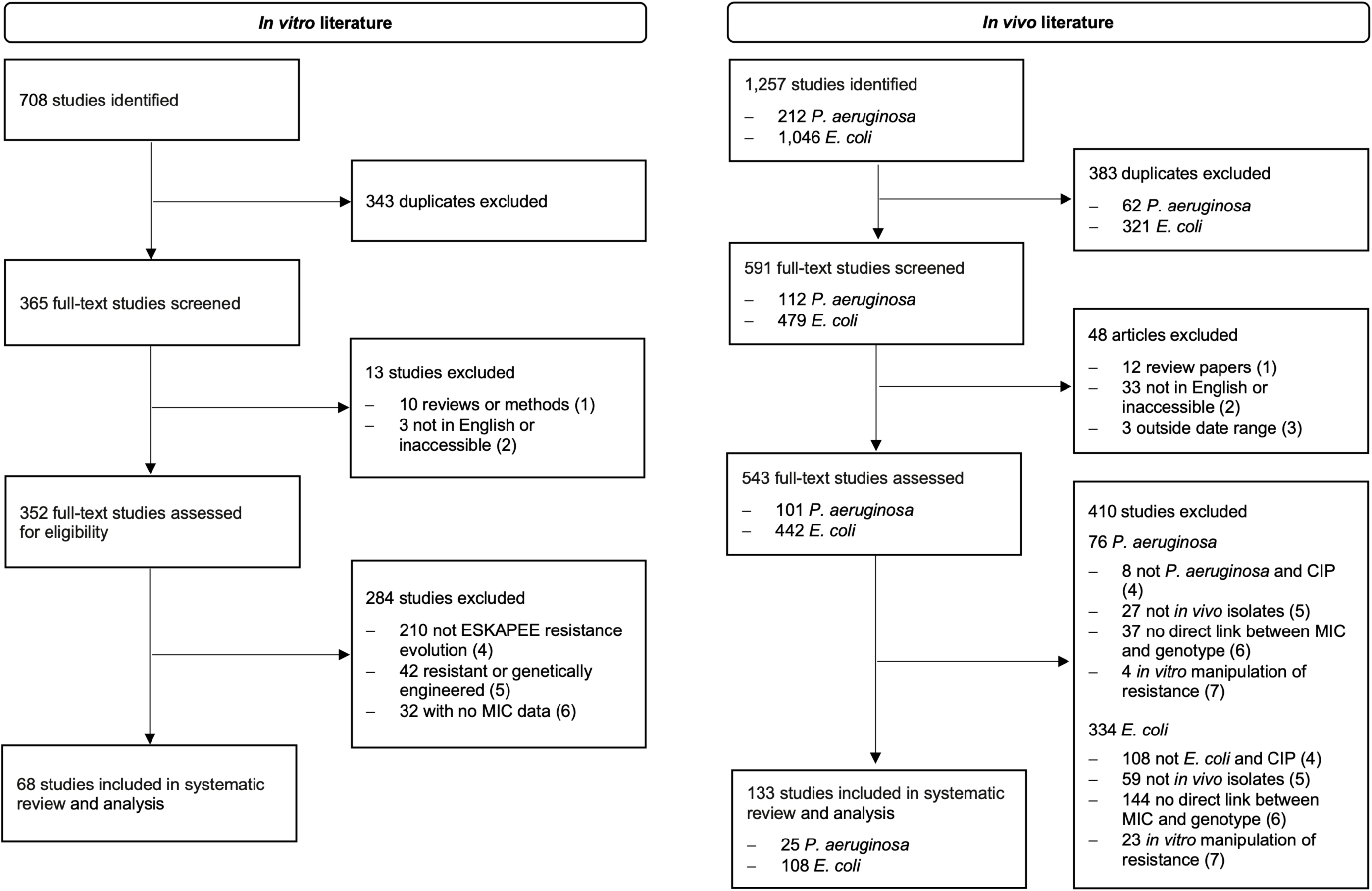
PRISMA detailing the workflow and numbers of publications found, screened and reviewed during this study.

**Fig. S2.**
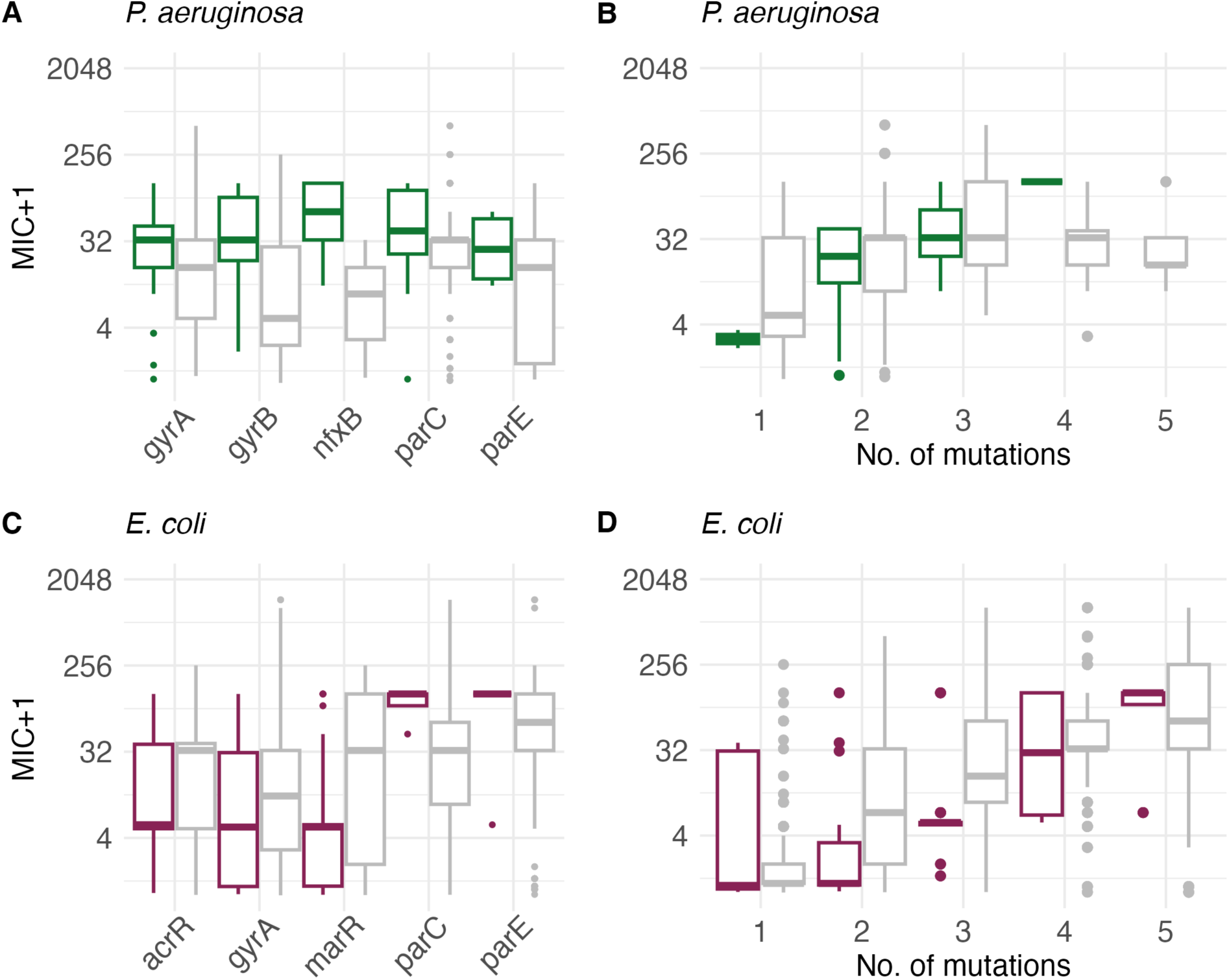
Evolved MIC by gene mutation and number of mutations in response to CIP exposure. The box plots illustrate the MIC values following exposure to CIP for the five most frequently selected gene mutations observed in resistant isolates of *P. aeruginosa* (A) and *E. coli* (C) across *in vitro* and *in vivo* studies. Furthermore, the MIC is presented alongside the corresponding number of mutations that emerged from CIP exposure in *P. aeruginosa* (B) and *E. coli* (D) isolates, both *in vitro* and *in vivo*. The figures present data restricted to isolates harbouring six or fewer mutations. MIC values are presented on a log2 scale. The colour scheme for the *in vitro* data of *P. aeruginosa* (green) and *E. coli* (burgundy red) has been retained. *In vivo* data are shown in grey.

**Fig. S3.**
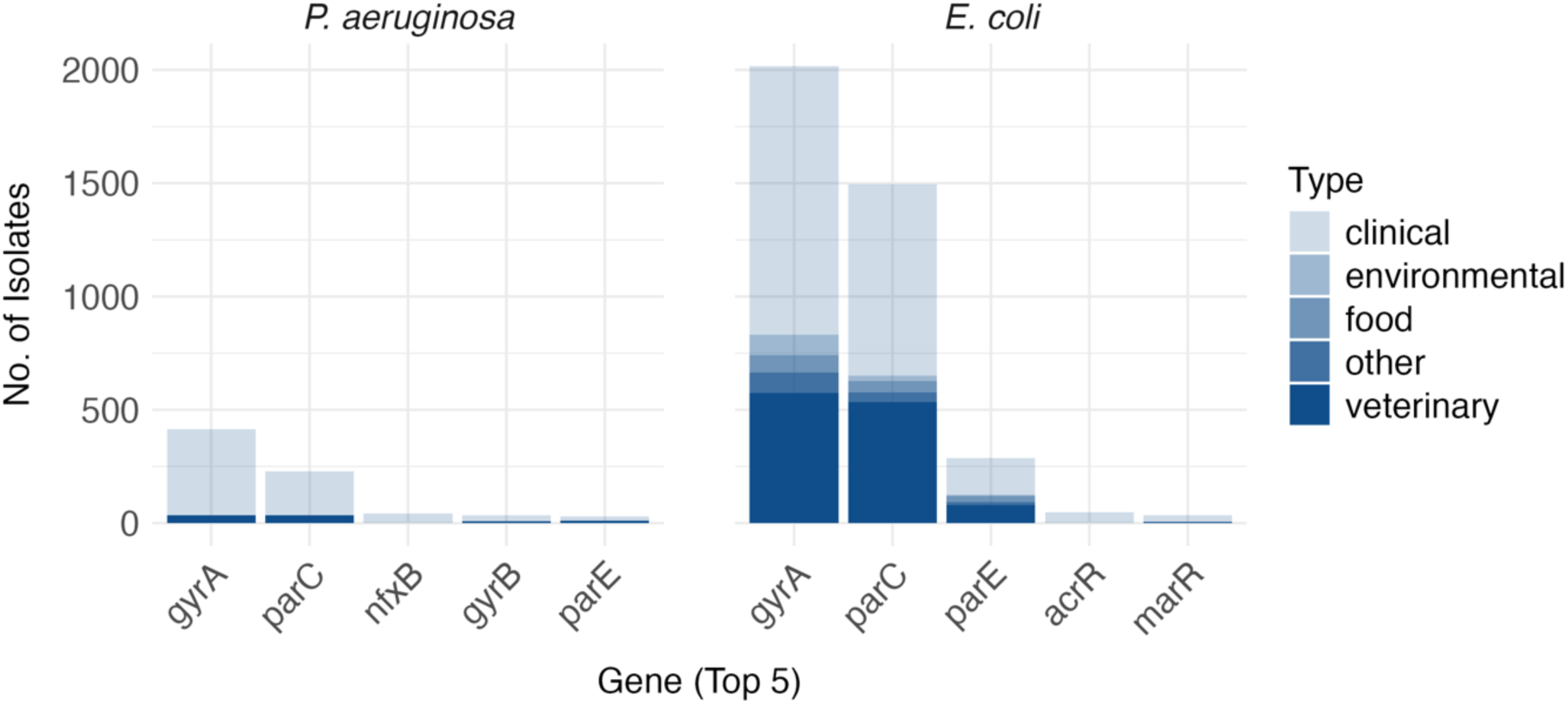
Sampling contexts among the *in vivo* isolates of *P. aeruginosa* and *E. coli*.

**Fig. S4.**
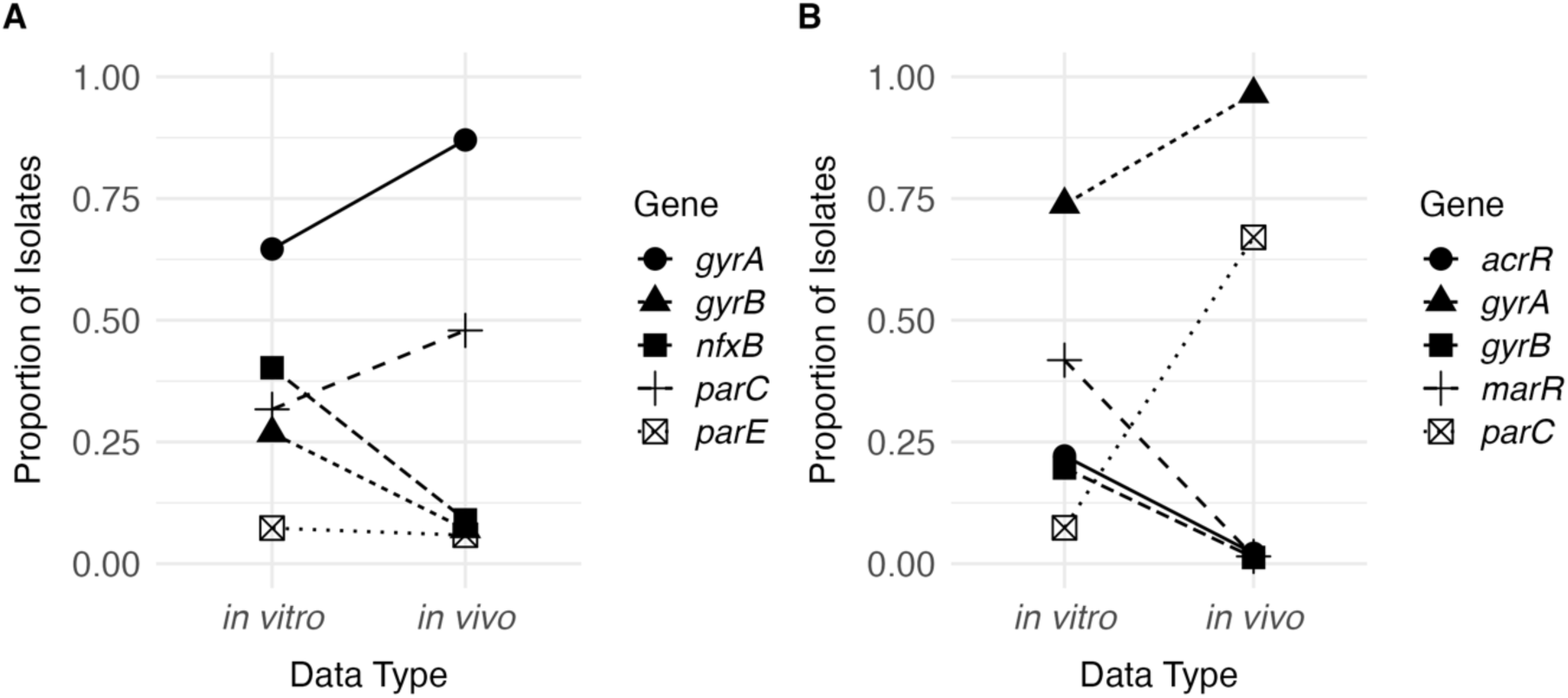
Proportions of detected genes following CIP exposure. Proportions of the top five detected gene mutations in *in vitro* and *in vivo* isolates of *P. aeruginosa* (A) and *E. coli* (B).

**Table S1.**
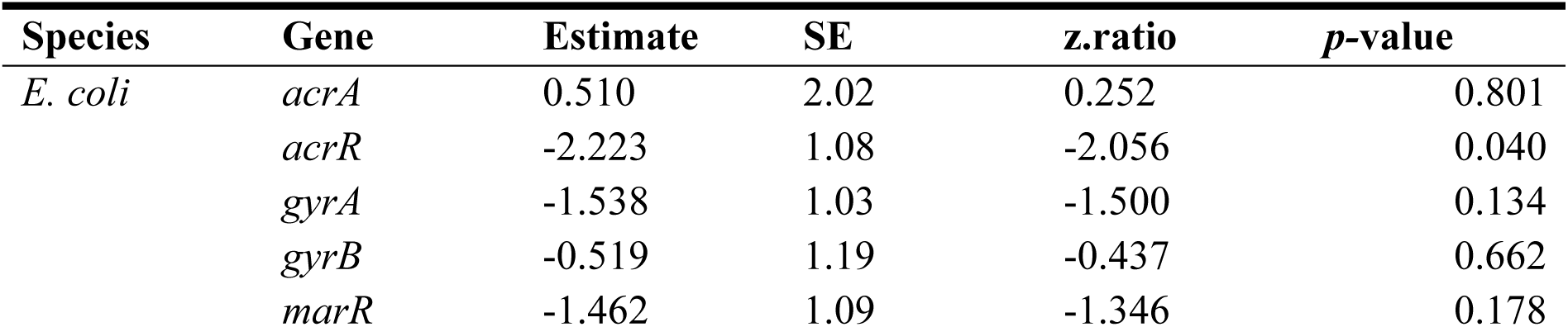
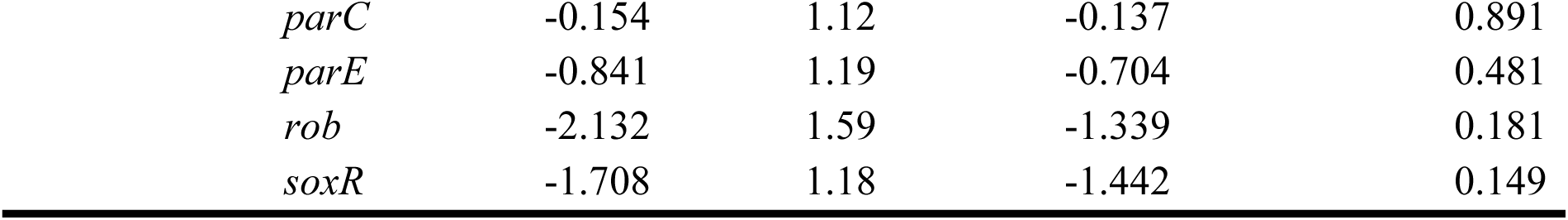
Pairwise comparison of *in vitro* and *in vivo* evolved MIC in *E. coli* isolates across gene mutations. The comparison was performed using Tukey’s post hoc test.

| Species | Gene | Estimate | SE | z.ratio | p-value |
| --- | --- | --- | --- | --- | --- |
| *E. coli* | *acrA* | 0.510 | 2.02 | 0.252 | 0.801 |
|  | *acrR* | -2.223 | 1.08 | -2.056 | 0.040 |
|  | *gyrA* | -1.538 | 1.03 | -1.500 | 0.134 |
|  | *gyrB* | -0.519 | 1.19 | -0.437 | 0.662 |
|  | *marR* | -1.462 | 1.09 | -1.346 | 0.178 |

|  |  |  |  |  |
| --- | --- | --- | --- | --- |
| *parC* | -0.154 | 1.12 | -0.137 | 0.891 |
| *parE* | -0.841 | 1.19 | -0.704 | 0.481 |
| *rob* | -2.132 | 1.59 | -1.339 | 0.181 |
| *soxR* | -1.708 | 1.18 | -1.442 | 0.149 |

